# Spinal rotational dynamics orchestrate locomotor recovery

**DOI:** 10.64898/2026.07.21.739844

**Authors:** Julian Colard, Jonathan Harnie, Rune W Berg, Alain Frigon

## Abstract

Locomotion is one of the most essential functions of the nervous system, yet the principles that generate it and how it might be restored after spinal cord injury remain unresolved. Walking is possible even after complete spinal cord injury and a traditional assumption is that this activity emerges from modular flexor-extensor transitions driven by somatosensory feedback. To test this assumption, we recorded bilateral motor unit activity from sixteen flexor/extensor hindlimb muscles in awake cats walking on a treadmill. We found that spinal motor populations operate according to a fundamentally different principle. Rather than alternating between flexion and extension, population activity continuously rotated through all phases of the locomotor cycle within a low-dimensional neural manifold. This rotational organisation persisted after complete spinal cord injury, revealing the preservation of a cyclic attractor despite substantial alterations in motor outputs. Computational modeling further indicated that locomotor activity was more consistent with an autonomous regime than with somatosensory-driven control. These findings suggest that locomotor recovery after complete spinal cord injury reflects the preservation of an intrinsic spinal attractor and that somatosensory feedback contributes to, rather than generates, the locomotor rhythm. More broadly, they establish rotational population dynamics as a fundamental principle of spinal locomotor recovery.

## MAIN

Walking is a fundamental motor behaviour that depends on the coordinated activity of distributed neural circuits spanning the brain and spinal cord. A complete thoracic spinal cord injury (SCI) permanently interrupts descending supraspinal projections and disrupts functional connectivity within lumbosacral spinal networks controlling locomotion of the legs/hindlimbs. This disconnection leads to an immediate loss of voluntary motor control and an initial phase of motor atonia, classically referred to as spinal shock^1^. At first glance, locomotor generation appears critically dependent on descending commands from the brain. However, decades of experimental work in mammalian models have demonstrated that spinal circuits retain an intrinsic capacity to generate rhythmic and coordinated locomotor activity even after complete transection^2,3^. Under such conditions, strengthening somatosensory afferent inputs promotes the re-emergence of functional locomotor patterns despite the persistent absence of supraspinal control. This capacity is generally attributed to the lumbosacral locomotor central pattern generator (CPG), which is traditionally thought to consist of distinct, reciprocally organized modules, in which individual neurons are specialized for either flexor or extensor activity, producing alternating rhythmic motor output^2–4^. However, recent studies challenge this classical binary and modular view of the CPG. Rather than arising from the alternation of discrete flexor–extensor modules, locomotor activity may instead emerge from interactions in the spinal network^5^. In this framework, rhythmic motor patterns are generated by low-dimensional population activity evolving continuously over time, reflecting intrinsic network properties rather than the activity of dedicated, functionally segregated neuronal groups.

In this way, locomotion in spinal-transected animals (i.e. spinal locomotion) is not merely the expression of an isolated oscillator. CPG activity is strongly shaped by proprioceptive and cutaneous afferent inputs, which regulate phase timing, burst amplitude, and intermuscular coordination^6–8^. Following spinal transection, dependence on somatosensory feedback becomes dominant, leading to conceptual frameworks in which locomotion is described as a state-dependent control system, where somatosensory inputs partially substitute for descending commands and increase their influence on phase transitions^9^. Although this view emphasises the functional role of somatosensory afferents, a central mechanistic question remains unresolved: is spinal locomotion primarily driven by afferent inputs, or does it emerge from the intrinsic structural and dynamical properties of the spinal network?

From this perspective, understanding spinal locomotor control requires access to neuronal population dynamics rather than to global EMG envelopes alone. In animal models, both before and after SCI, locomotor control has been extensively investigated at the levels of the spinal circuitry and whole muscle output, including in lamprey and fish^10,11^, in rodents^4,12^, and in mammalian models, such as the cat ^2^, including air-stepping paradigms, acute preparation^13^. However, analyses of spinal locomotor activity at the motoneuronal population level during awake, load-bearing real locomotion, particularly in chronic conditions, remain comparatively limited. To our knowledge, this study provides the first such characterization across the transition from intact to chronic complete spinal cord injury. Recording simultaneously from multiple motor units across different muscle pools provides a valuable window into collective dynamics, as motor unit activity enables the decoding of the underlying spinal motoneuron population dynamics^14^. Moreover, recent work has demonstrated a tight and temporally precise relationship between spinal interneuron population activity and muscle output, with high predictive power on millisecond timescales^13^. Together, these findings confirm the idea that motor unit recordings capture a meaningful projection of the underlying spinal network dynamics, including coordinated activity across interneuron and motoneuron populations.

In this context, we sought to determine whether spinal locomotor organisation is primarily driven by external inputs or instead emerges from the intrinsic collective dynamics of the spinal network. To address this, we characterised motoneuronal population activity during locomotion using a state-space approach, enabling comparison of the organisation and evolution of population dynamics before and after spinal transection. By directly accessing the collective structure of motoneuronal activity, this framework moves beyond descriptions based solely on muscle output and reveals how neuronal states unfold, organise, and evolve throughout the locomotor cycle. We therefore examined how the shape, stability, and temporal organisation of these dynamics are affected by the loss of descending inputs, and to what extent their fundamental properties are preserved or altered.

We hypothesized that locomotion following spinal transection is governed by an intrinsic low-dimensional collective dynamic, the global structure of which persists despite the loss of supraspinal control. In this framework, somatosensory afferents do not constitute the primary driver of locomotor activity, but instead contribute to the initiation of the dynamics, allowing the network to enter a self-sustained regime.

To test this hypothesis, we combined intramuscular recordings of motor unit activity with a recurrent dynamical model where we selectively manipulated different input sources to assess their contribution to the observed population dynamics and to determine whether these dynamics can emerge independently of external inputs.

### Rotational spinal motoneuron activity during locomotion

Chronic intramuscular EMG electrodes implanted bilaterally in eight flexor and eight extensor hindlimb muscles enabled simultaneous monitoring of muscle-level and motor unit activity during treadmill locomotion in 15 cats **(Fig. 1A)**. Following a complete thoracic transection, recordings were obtained in same cats to isolate the intrinsic contribution of spinal circuits in the absence of supraspinal input **(Fig. 1B)**. At the level of muscle activity, locomotion was characterised by the expected alternation between flexor and extensor bursts **(Fig. 1C)**, a pattern that persisted after transection **(Fig. 1D)**. This preservation suggests that the rhythmic structure remains embedded within the spinal circuitry despite the absence of supraspinal drive. At the level of individual motor units, population activity initially appeared disorganised. Spike rasters revealed heterogeneous and temporally dispersed discharge patterns, with no readily apparent structure when viewed directly in intact **(Fig. 1C)** or transected **(Fig. 1D)** conditions. This apparent lack of organisation reflects the complexity of population activity when visualised in the space of individual units. To reveal potential structure, we reordered motor units according to their preferred discharge phase within the locomotor cycle. This simple transformation revealed a striking organisation, with motor unit activity forming a continuous diagonal sequence spanning the entire cycle **(Fig. 1E)**. Remarkably, this continuous organisation persisted after spinal transection **(Fig. 1F)**, although it appeared less sharply defined. The preservation of this phase-ordered sequence despite the absence of descending input suggests that it reflects an intrinsic property of the spinal circuitry rather than a supraspinal imposed control. When pooling neurons across trials and animals (n = 15), preferred phases covered the entire locomotor cycle in both intact and transected conditions (Fig. 1G, H), revealing a continuous progression of activity across the cycle rather than segregation into discrete flexor- and extensor-related clusters. This coverage was continuous but not flat: cumulative phase distributions departed from perfect circular uniformity (Kuiper test). A group-level comparison against uniform surrogate datasets confirmed that these deviations exceeded those expected by chance. Such organisation is characteristic of rotational population neuronal dynamics, previously described in turtle movements^5^.

**Figure 1.**
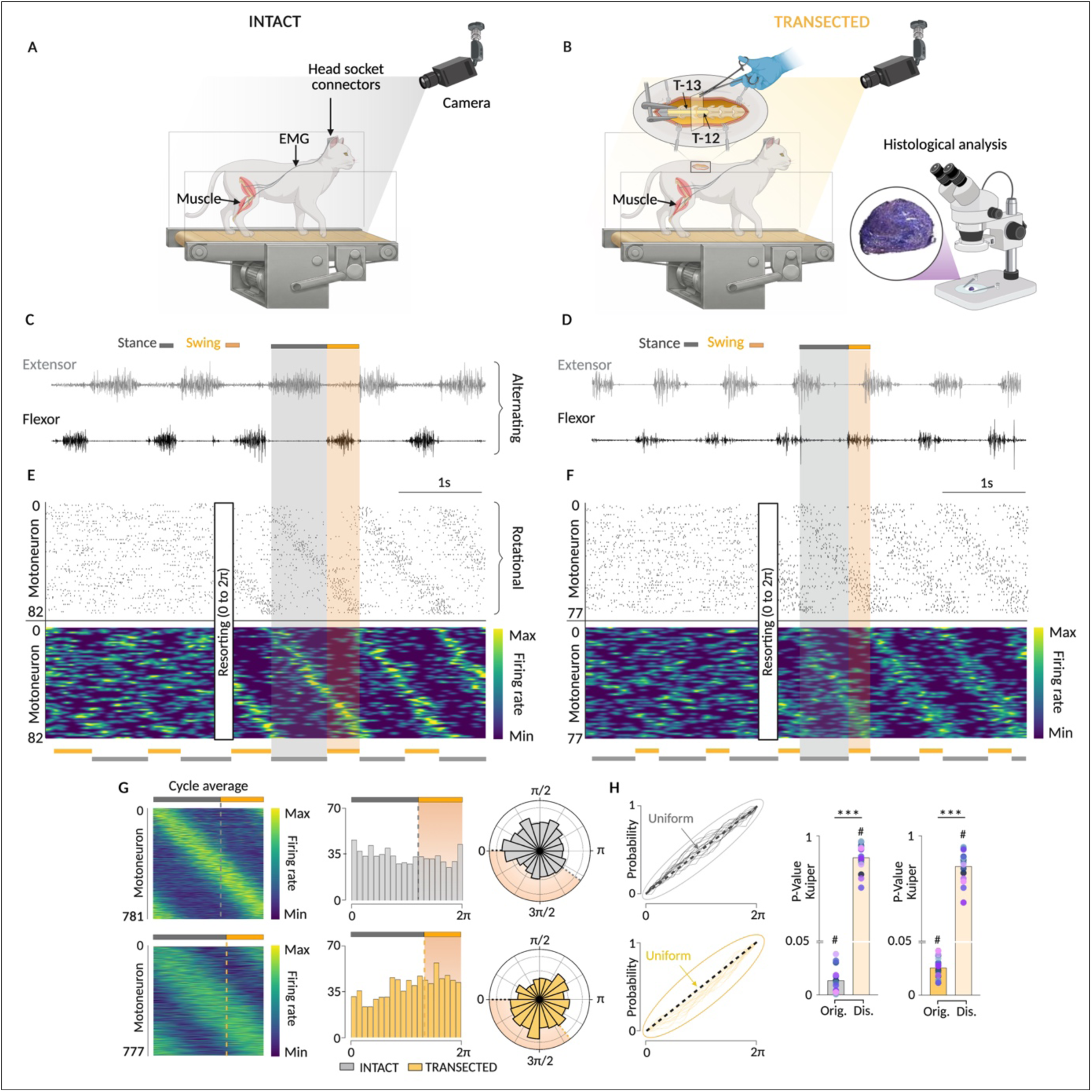
Rotational population dynamics of spinal motoneurons during walking are preserved after spinal transection. **A)** Experimental setup in intact cats. Intramuscular EMG recordings were obtained from 16 hindlimb muscles during treadmill locomotion. High-speed videography was used to identify stance (grey) and swing (orange) phases. **B)** Experimental setup 6–8 weeks after complete spinal transection at T12–T13. Recording conditions, including treadmill speed (0.4 m s⁻¹), were matched to the intact condition. Histology confirmed complete transection. **C)** Representative raw EMG recordings from vastus lateralis (top) and anterior sartorius (bottom) during intact locomotion. **D)** Representative raw EMG recordings from soleus (top) and tibialis anterior (bottom) after spinal transection. **E–F)** Motoneuron population activity sorted by locomotor phase. Spike rasters (top) and firing-rate estimates (bottom) are ordered according to the circular phase of peak discharge (0–2π) in intact (E; N = 82 motor units) and transected (F; N = 77 motor units) conditions. **G)** Phase-sorted population activity pooled across animals (n = 15; 781 motor units in intact and 777 in transected conditions), shown in linear and circular representations. **H)** Cumulative phase distributions for intact (top) and transected (bottom) conditions. Deviations from circular uniformity were assessed using the Kuiper test (#, P < 0.001). Group-level significance was confirmed by comparison with surrogate distributions using a Wilcoxon rank-sum test (***, P < 0.001). EMG, electromyography.

To quantify how spinal transection alters motor output, we next examined motor unit discharge properties across locomotor phases using motor units identified across 16 hindlimb muscles **(Fig. 2A)**. Transection led to a marked reduction in discharge rates during stance but not during swing phase **(Fig. 2B)**, with the strongest effects observed in extensors **(Supplementary Table 1)**. In contrast, the temporal structure of motor unit activity was largely preserved, as discharge duration windows showed only a modest effect, restricted to flexors during swing **(Fig. 2C and Supplementary Table 2)**. Transection markedly reduced discharge rate but affected discharge duration to a proportionally much smaller extent, indicating that descending pathways act primarily on the intensity of motoneuronal activity, whose temporal organisation remains largely determined by spinal mechanisms.

**Figure 2.**
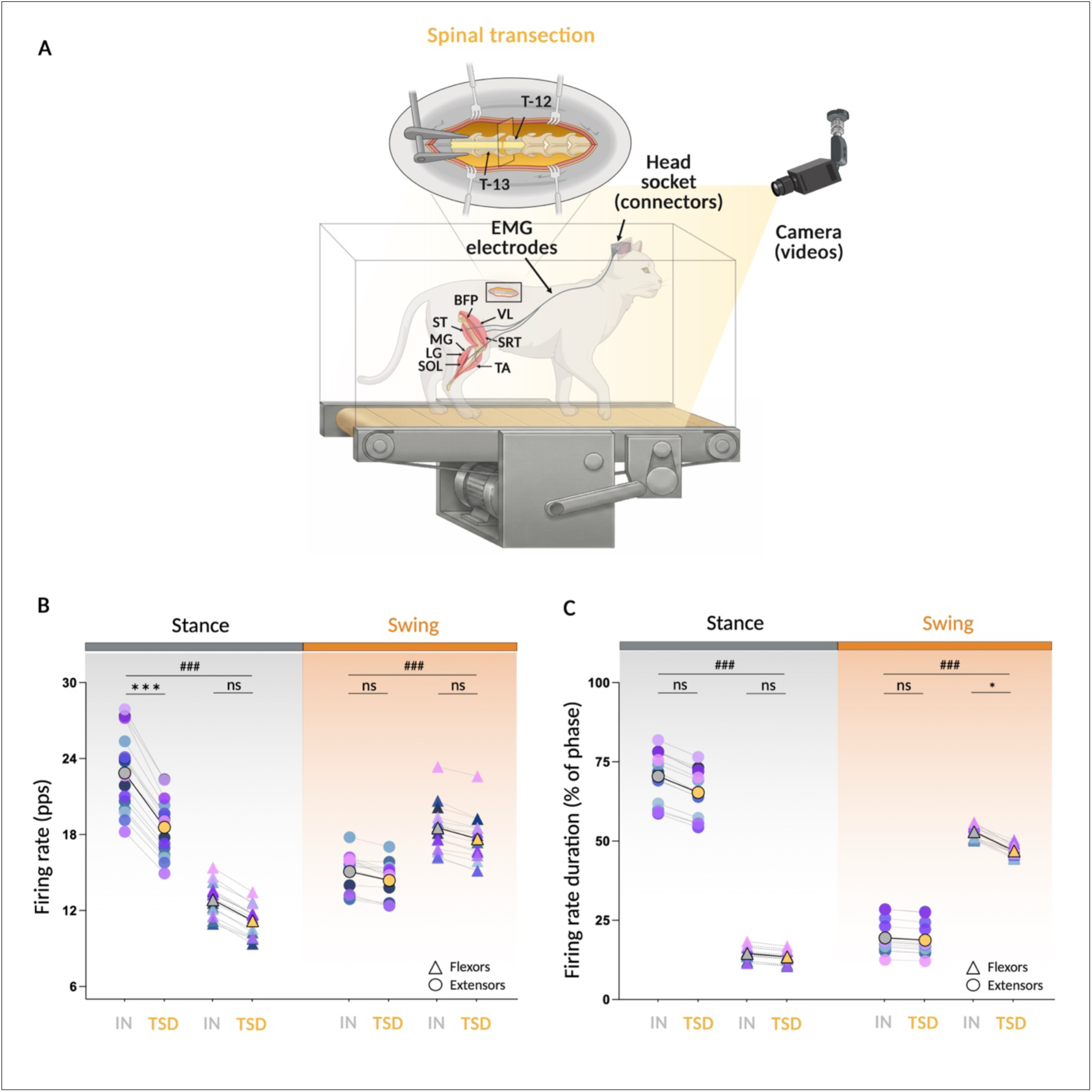
Motor unit characteristics before and after spinal transection. Data were obtained from 15 cats in the intact (IN) and transected (TSD) conditions. **A)** Schematic illustration of the experimental design for motor unit recordings from 16 hindlimb muscles (8 muscles recorded bilaterally). **B)** Mean motor unit discharge rate during stance (grey) and swing (orange). **C)** Mean discharge duration expressed as a percentage of the locomotor phase during stance (left) and swing (right). Each point represents one animal; colours identify individual cats, and black lines indicate paired comparisons. Motor units were grouped according to function as extensors (circles; LG, lateral gastrocnemius; MG, medial gastrocnemius; SOL, soleus; VL, vastus lateralis) or flexors (triangles; BFP, biceps femoris posterior; ST, semitendinosus; SRT, anterior sartorius; TA, tibialis anterior). Statistics: ns, not significant; *, P < 0.05; ***, P < 0.001 for intact vs transected. ###, P < 0.001 for extensors vs. flexors. Mn, motoneuron.

### Low-dimensional motoneuron population dynamics

To decode how spinal transection affects the functional strategy orchestrating motor output, we must explore the low-dimensional collective dynamics and the geometry of the state space. Projection onto the first principal components revealed that population activity evolves along a structured, low-dimensional trajectory **(Fig. 3A, top)**. In intact animals, trajectories formed a compact and highly reproducible cycle, with a clear progression through the stance and swing phases. Topological analysis identified a dominant homology H_1_ component in the absence of higher-order features, consistent with a ring-shaped manifold indicating cyclic population dynamics. Following spinal transection, this cyclic structure persisted but was less sharply defined. Trajectories were more dispersed and less tightly organised **(Fig. 3A, bottom)**, and the persistence of the H_1_ component appeared less pronounced, suggesting a weakening of the underlying cyclic dynamics. The cumulative variance explained by principal components revealed a difference in the dimensional structure of population activity **(Supplementary Table 3)**. In intact animals, variance accumulated gradually across components, with 5 dimensions required to explain 80% of the variance, indicating that activity is distributed across multiple coordinated modes **(Fig. 3B, top)**. After spinal transection **(Fig. 3B, bottom)**, variance saturated more rapidly, with 3 dimensions sufficient to explain the same proportion of variance, indicating that activity is dominated by a reduced number of modes, consistent with the loss of descending inputs. Canonical correlation revealed a partial correspondence between intact and transected population activity **(Fig. 3C)**. The leading (first five) dimensions were correlated (r = 0.66), with significant correlations extending across multiple components, indicating that the underlying low-dimensional subspace is partially conserved after transection, although alignment remains incomplete.

**Figure 3.**
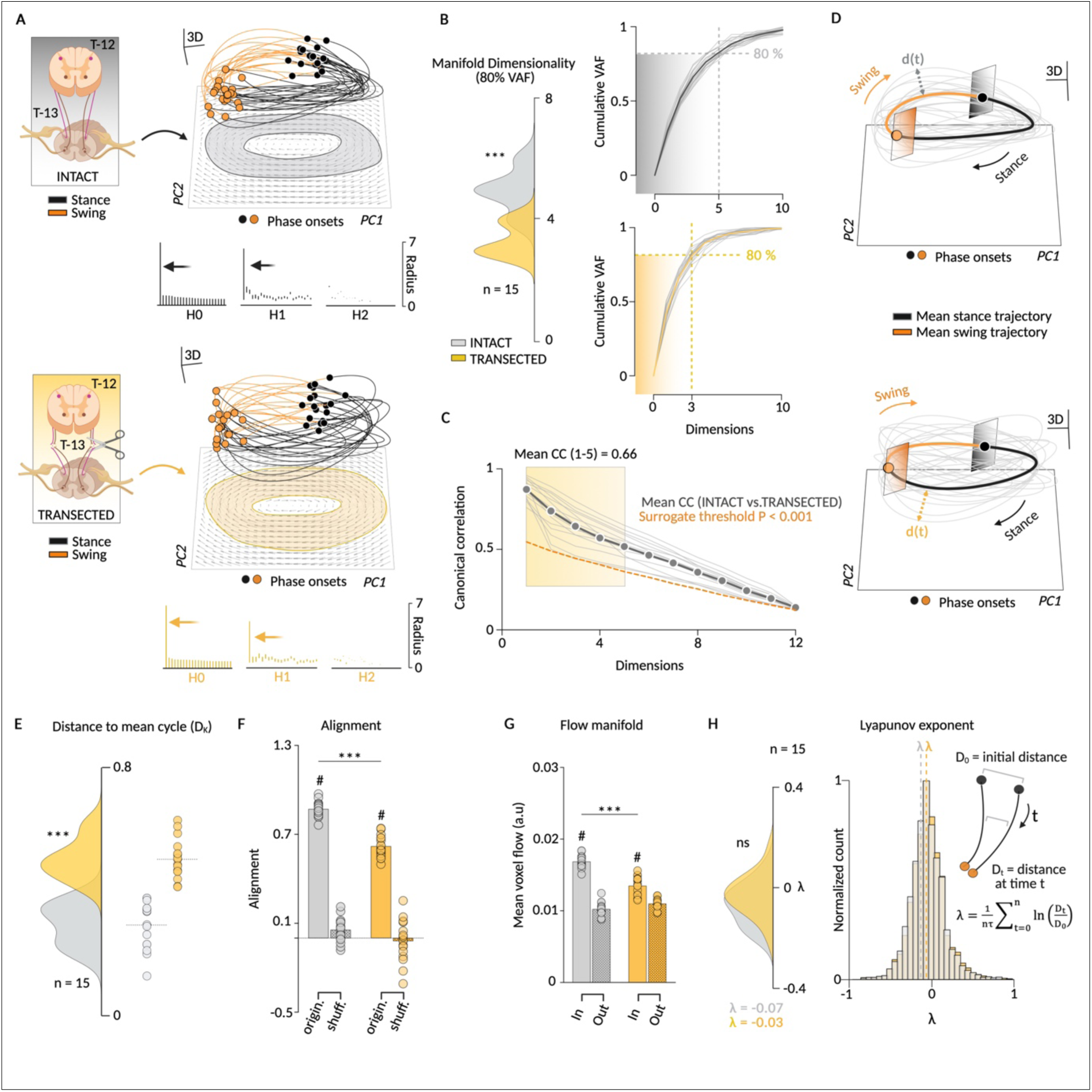
Spinal transection alters the collective dynamics and state-space geometry of spinal motoneurons. **A)** Experimental schematics for intact (top) and transected (bottom; T12–T13 lesion) conditions. Neural trajectories are shown in three-dimensional principal component (PC) space and as two-dimensional PC1–PC2 projections. Locomotor phase is colour-coded (stance, black; swing, orange), with dots indicating phase transitions. Arrows represent the state-space flow field. Persistent homology barcodes are shown below. **B)** Motoneuronal manifold dimensionality. Left, violin plots showing the number of dimensions required to explain 80% of the variance accounted for (VAF) in intact (grey) and transected (yellow) conditions. Right, cumulative VAF across latent dimensions. **C)** Canonical correlation analysis comparing intact and transected manifolds across 12 dimensions. Grey dots represent mean canonical correlation coefficients; the dashed orange line indicates the surrogate significance threshold (P < 0.001). **D)** Mean locomotor trajectories in three-dimensional PC space. Grey traces represent individual locomotor cycles. **E)** Mean distance of individual trajectories from the average locomotor cycle. **F)** Alignment of individual trajectories to the mean flow within the locomotor manifold compared with shuffled datasets. **G)** Mean voxel flow magnitude (a.u.) measured inside (In) and outside (Out) the manifold. **H)** Violin plots and histograms of Lyapunov exponents (λ) derived from local trajectory divergence in state space. ***, P < 0.001 for intact vs transected; #, P < 0.001 for Original vs. Shuffled and In vs. Out; ns, not significant.

Dimensionality alone, however, does not capture how population activity evolves within this preserved manifold. We therefore next examined the stability of trajectories within the manifold. In intact animals, state trajectories remained tightly clustered around the mean cycle (**Fig. 3D top)**, indicating high orbital stability and a reproducible progression through the latent space. Following transection **(Fig. 3D bottom)**, trajectories became more dispersed **(Supplementary Table 4)**. This increase in transverse variance relative to the mean cycle demonstrates that the system’s stability is reduced, resulting in heightened cycle-to-cycle variability **(Fig. 3E)**. Motoneuronal activity was strongly aligned with the flow field, far above its shuffled baseline **(Fig. 3F)**, consistent with a coherent evolution along the manifold. After transection this alignment was reduced, and the difference from the shuffled baseline narrowed **(Supplementary Table 5)**, indicating that trajectories deviated from the dominant flow and became less distinguishable from temporally unstructured activity. Flow was stronger within the manifold than outside **(Fig. 3G)** and weakened after transection specifically within the manifold, leaving off-manifold flow unchanged **(Supplementary Table 6)**. This selective loss reduced the contrast between the manifold and the surrounding state space, reflecting a degradation of the structured flow that guides the population dynamics. The Lyapunov exponent (λ) remained close to zero in both conditions **(Fig. 3H and Supplementary Table 7)**. Because this characterises a system in a steady-state regime, the result is consistent with the persistence of dynamics consistent with a limit-cycle attractor, despite the absence of descending inputs. In conclusion, while transection disrupts locomotor precision, the population activity continues to evolve along a cyclic trajectory, demonstrating the robustness of an intrinsic spinal attractor. Although its stability is diminished, with trajectories diverging more from the cycle and being less effectively guided by the flow the fundamental limit-cycle structure is preserved.

### Spinal locomotion theory

Having demonstrated that spinal motoneuron populations retain low-dimensional rotational dynamics even in the absence of supraspinal inputs, we now revisit the origin of this rhythmic activity. Recovery of locomotion following a spinal transection has largely been attributed to the driving role of somatosensory afferents. Here we propose instead that this dynamical attractor is not generated by somatosensory feedback but emerges intrinsically from the recurrent connections of the spinal cord. This architecture implements a state-dependent balance between excitation and inhibition and operates near a dynamical instability, giving rise to self-sustained population dynamics. The recurrent connectivity thus defines a low-dimensional dynamical space and shapes the flow within it, such that the system naturally follows structured trajectories. In this framework, motor output arises from intrinsically generated collective dynamics evolving along a low-dimensional manifold. Locomotion is therefore not driven by external inputs but emerges from the internal organisation of the network, which constrains and guides the evolution of activity. Somatosensory feedback does not generate these dynamics but can shift the system into a self-sustained regime. The rhythm therefore does not arise from any single component of the network but from the closed recurrent loop as a whole: attenuating the motoneuronal segment abolished cyclic structure even though the premotor recurrent update remained intact, indicating that the loop must close through the output population for self-sustained dynamics to emerge. Within this framework, the motoneuronal population is not external to the rhythm-generating network but part of its recurrent architecture, consistent with anatomical evidence for recurrent motoneuron collaterals and Renshaw-mediated feedback within the spinal cord.

To test this theory, we developed animal-specific latent recurrent dynamical models trained individually on multi-muscle motoneuronal populations recorded during intact locomotion. The network integrates two external inputs (i.e., descending supraspinal drive and phase-dependent somatosensory feedback) alongside an explicit, low-dimensional representation of the network’s collective state, derived from the population output at the preceding time step, isolable and manipulable in silico. Recurrent units do not correspond to identified spinal neurons or cell classes. They instead provide a computational substrate for the collective dynamics generated by the spinal locomotor network. Motor unit activity is not represented as individual units within the recurrent network, but a low-dimensional summary of the population output is fed back into the recurrent update, such that the motoneuronal population both reads out and participates in the collective dynamics. The CPG thus emerges from the network’s intrinsic dynamics rather than being implemented as an explicit flexion– extension oscillator. Selectively ablating these inputs in silico and regenerating network activity in a purely autoregressive, closed-loop manner (without access to ground-truth activity beyond a brief initialisation period) formally disentangled their relative contributions to the locomotor manifold **(Fig. 4)**. Projecting population activity onto the first three principal components revealed that the compact manifold of the biological baseline **(Fig. 4A, left)** was faithfully mirrored by the intact in silico model **(Fig. 4A, second panel)**. Persistent homology analysis confirmed this shared topology, with a robust H_1_ component detected in both biological and in silico intact conditions, consistent with a stable ring-like structure of the locomotor attractor. This topological signature was preserved in the in silico spinal-transected configuration, with more elongated trajectories and reduced H_1_ persistence, consistent with a weakened cyclic structure **(Fig. 4A, third panel)**. In contrast, with somatosensory afferent-only drive, the H_1_ component was absent, indicating a loss of cyclic topology **(Fig. 4A right)**. The cumulative VAF revealed a progressive reduction in dimensionality across conditions **(Fig. 4B)**, with both biological and in silico intact configurations requiring five components to explain 80% of the variance **(Fig. 4B, left and second panels)**, compared to three in the in silico spinal-transected condition **(Fig. 4B, third panel)** and one under somatosensory-only drive **(Fig. 4B, right)**. These differences were confirmed at the group level **(Fig. 4C)**, with no significant difference between biological and in silico intact conditions, but a marked reduction in the in silico spinal-transected configuration, further accentuated under somatosensory-only drive **(Supplementary Table 8)**. Consistent with this, canonical correlation analysis relative to the biological intact condition revealed strong alignment of low-dimensional subspaces in the in silico intact model **(Fig. 4D)**, which remained partially preserved for in silico spinal-transected, but was lost under somatosensory-only drive, indicating a breakdown of the shared subspace. The flow within the manifold was well organised in the in silico intact model **(Fig. 4E–F)**, with high alignment **(Supplementary Table 9)** and stronger flow inside than outside the manifold **(Supplementary Table 10)**, consistent with coherent limit cycle attractor dynamics. This organisation remained partially preserved after transection, with trajectories still largely following the flow, although less strongly. In contrast, under somatosensory-only drive, both alignment and flow collapsed, indicating a loss of organised dynamics. Together, these results suggest that somatosensory feedback alone is insufficient to sustain locomotor dynamics and that these dynamics likely rely on intrinsic network mechanisms preserved after transection.

**Figure 4.**
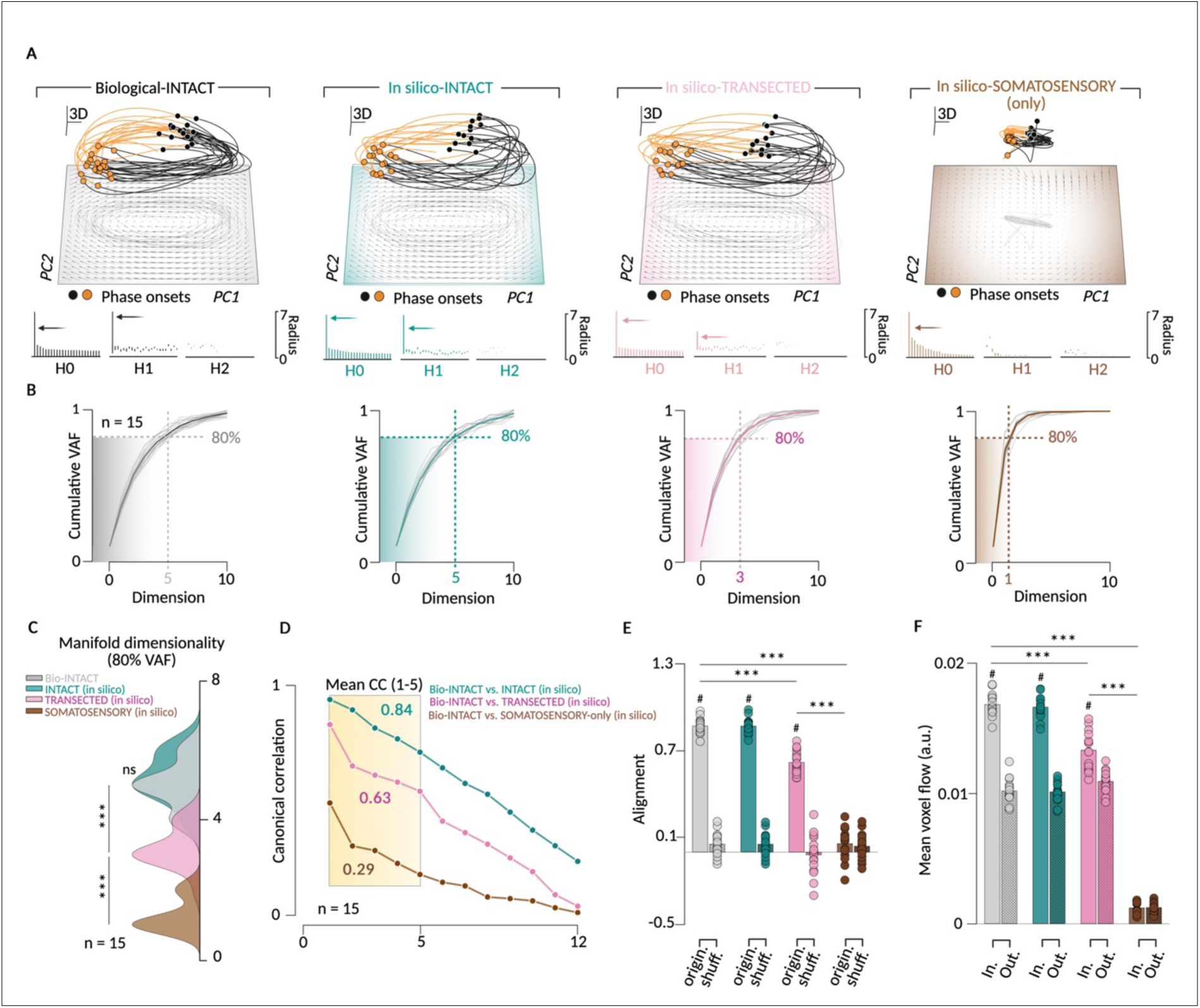
Somatosensory feedback alone is insufficient to sustain the cyclic structure of population activity. **A)** Motoneuronal trajectories in three-dimensional principal component (PC) space (top) and corresponding PC1– PC2 projections (bottom) for biological data (Biological-INTACT) and in silico models (INTACT, TRANSECTED, and SOMATOSENSORY-only). Locomotor phase is colour-coded (stance, black; swing, orange), with dots indicating phase transitions. Arrows represent the state-space flow field. Persistent homology barcodes are shown below. **B)** Cumulative variance accounted for (VAF) of motoneuronal activity for biological data (grey) and in silico models (cyan, pink, and brown; n = 15). **C)** Violin plots showing the number of dimensions required to explain 80% of the variance in biological and in silico conditions (n = 15). **D)** Canonical correlation analysis comparing low-dimensional manifolds between the biological intact condition and in silico models across 12 dimensions (n = 15). **E)** Alignment between locomotor trajectories and the estimated state-space flow field for biological data and in silico models. Bars indicate original and shuffled datasets. **F)** Mean voxel flow magnitude measured inside (In) and outside (Out) the manifold for biological data and in silico models. ***, P < 0.001 for intact vs transected; #, P < 0.001 for Original vs. Shuffled and In vs. Out; ns, not significant.

### Locomotion as an emergent property of spinal population dynamics

We next examined whether spinal locomotor population dynamics observed in recordings could be reproduced by the trained recurrent neural network (RNN) under simulated transection conditions **(Fig. 5)**. To do so, descending supraspinal inputs were removed while somatosensory feedback was preserved. We then systematically varied the gain applied to the collective-state feedback signal, which conveys a low-dimensional representation of the network’s own previous output back into the recurrent update. Reducing this gain progressively limited the influence of the network’s past activity on future states, while leaving the somatosensory input and the recurrent update itself unchanged. As this feedback was attenuated, the topology of the resulting trajectories was progressively altered **(Fig. 5A)**. When it was fully suppressed (0.00), trajectories became confined to a low-dimensional subspace and exhibited back-and-forth dynamics rather than forming a closed loop **(Fig. 5A, top right)**. Consistently, persistent homology no longer detected an H1 component, indicating a loss of cyclic topology. Notably, this loss of cyclic structure occurred despite the presence of phase-dependent somatosensory input and of an intact recurrent update, indicating that phase information alone is not sufficient to generate cyclic population dynamics in the absence of collective-state feedback. Increasing the gain (0.50) restored an H1 component, indicating the re-emergence of a cyclic manifold and recapitulating the ring-like topology observed with the biological transection. Notably, population activity remained organized around a closed orbit, consistent with the persistence of a limit-cycle attractor despite the loss of descending supraspinal drive **(Fig. 5A, bottom right)**.

**Figure 5.**
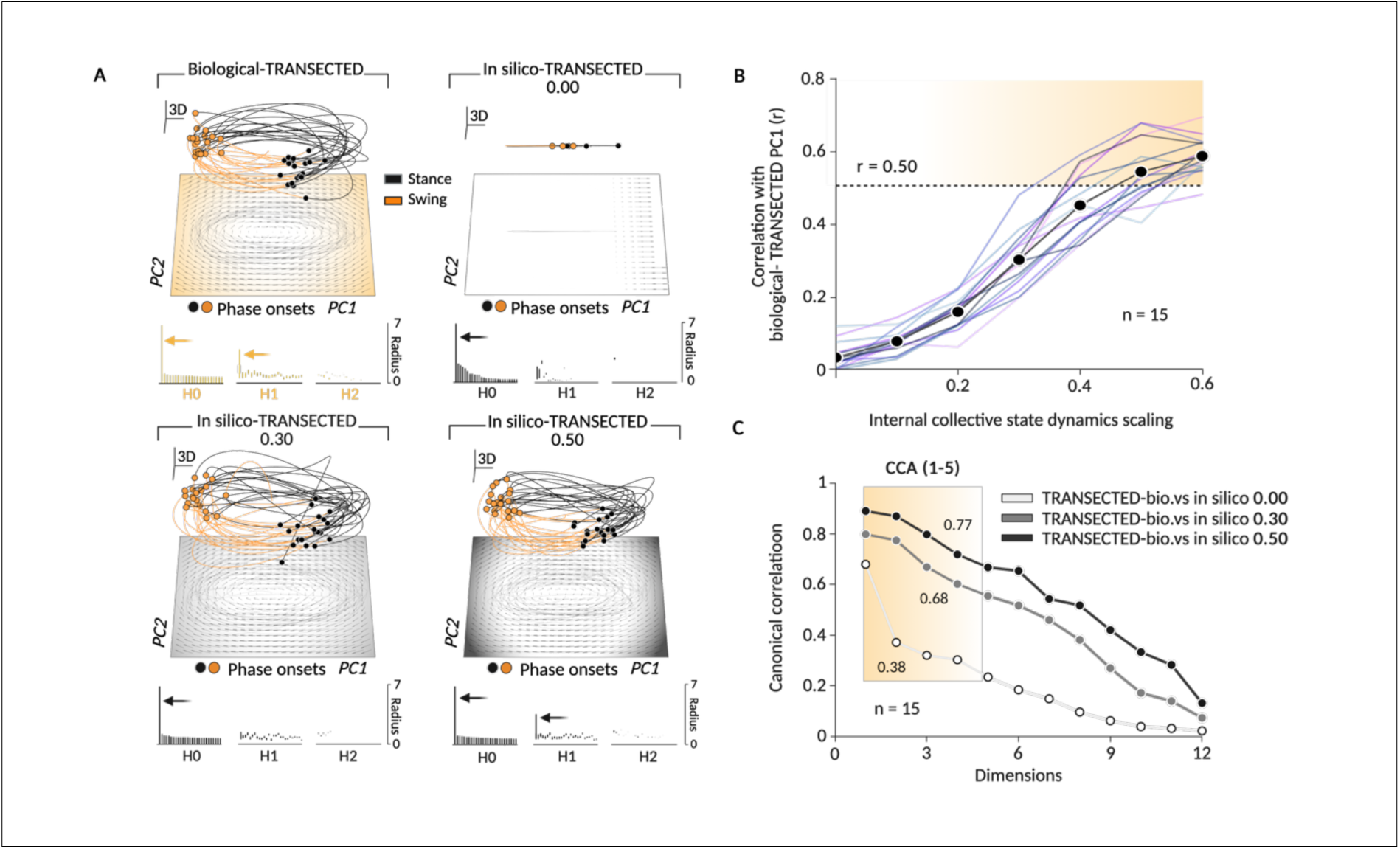
Spinal locomotion emerges from self-sustained network dynamics. **A)** Motoneuronal trajectories in three-dimensional principal component (PC) space (top) and corresponding PC1–PC2 projections (bottom) for biological data (Biological-TRANSECTED) and in silico models at three representative different collective-state feedback gains (TRANSECTED 0.00, 0.30, and 0.50). Locomotor phase is colour-coded (stance, black; swing, orange), with dots indicating phase transitions. Arrows represent the state-space flow field. **B)** Correlation between biological and in silico PC1 trajectories as a function of the collective-state feedback gain (n = 15). The black curve indicates the population mean. **C)** Canonical correlation analysis across 12 dimensions comparing biological data and in silico models with different collective-state feedback gains (n = 15). Values within the shaded region indicate the mean canonical correlation across the first five dimensions (CC1–5).

To quantify similarity, we computed the correlation between the first principal component (PC1) of the biological transected data and that of the in silico model across scaling values **(Fig. 5B)**. Correlation increased monotonically with collective-state feedback gain, with a scaling value of 0.50 required to reach a correlation of r = 0.50 with the biological reference. Canonical correlation analysis further revealed that subspace alignment increased with scaling, with higher scaling conditions showing strong alignment across the leading modes, while correlations decreased with mode order, indicating that shared structure is concentrated in low-dimensional components (**Fig. 5C).**

We next used the trajectory tangling metric to assess how reliably future population activity can be inferred from the current state. Tangling quantifies the divergence of nearby trajectories over time and thus distinguishes internally driven from input-driven regimes. In the absence of collective-state feedback (gain = 0), tangling values were markedly higher than those observed in biological transected data, indicating that population evolution was weakly constrained by the current state and more strongly influenced by inputs **(Fig. 6A)**. As collective-state feedback was progressively restored, this discrepancy diminished, with tangling values converging towards the biological range. Intermediate and higher scaling conditions fell within a regime consistent with locally predictable dynamics. Consistently, spinal population states revealed that local predictability increased with collective-state feedback gain, with tangling values decreasing towards a reference range defined by autonomous RNN dynamics **(Fig. 6B)**. This convergence held in every animal taken individually **(Fig. 6C and Supplementary Table 11)**. At zero gain, tangling was higher than in the biological transected condition in all 15 animals, with near-complete separation of the distributions and the effect remained significant in all 15 animals at a gain of 0.10. It weakened at 0.30, where 11 of 15 animals still differed from the biological condition, and largely vanished at 0.50, where only 3 of 15 did. The median effect size therefore decreased monotonically with feedback gain, from complete separation to differences that were no longer detectable. Together, these results suggest that, following spinal transection, somatosensory inputs influence population activity but do not primarily drive its evolution. Instead, population dynamics may be predominantly governed by intrinsic collective state dynamics.

**Figure 6.**
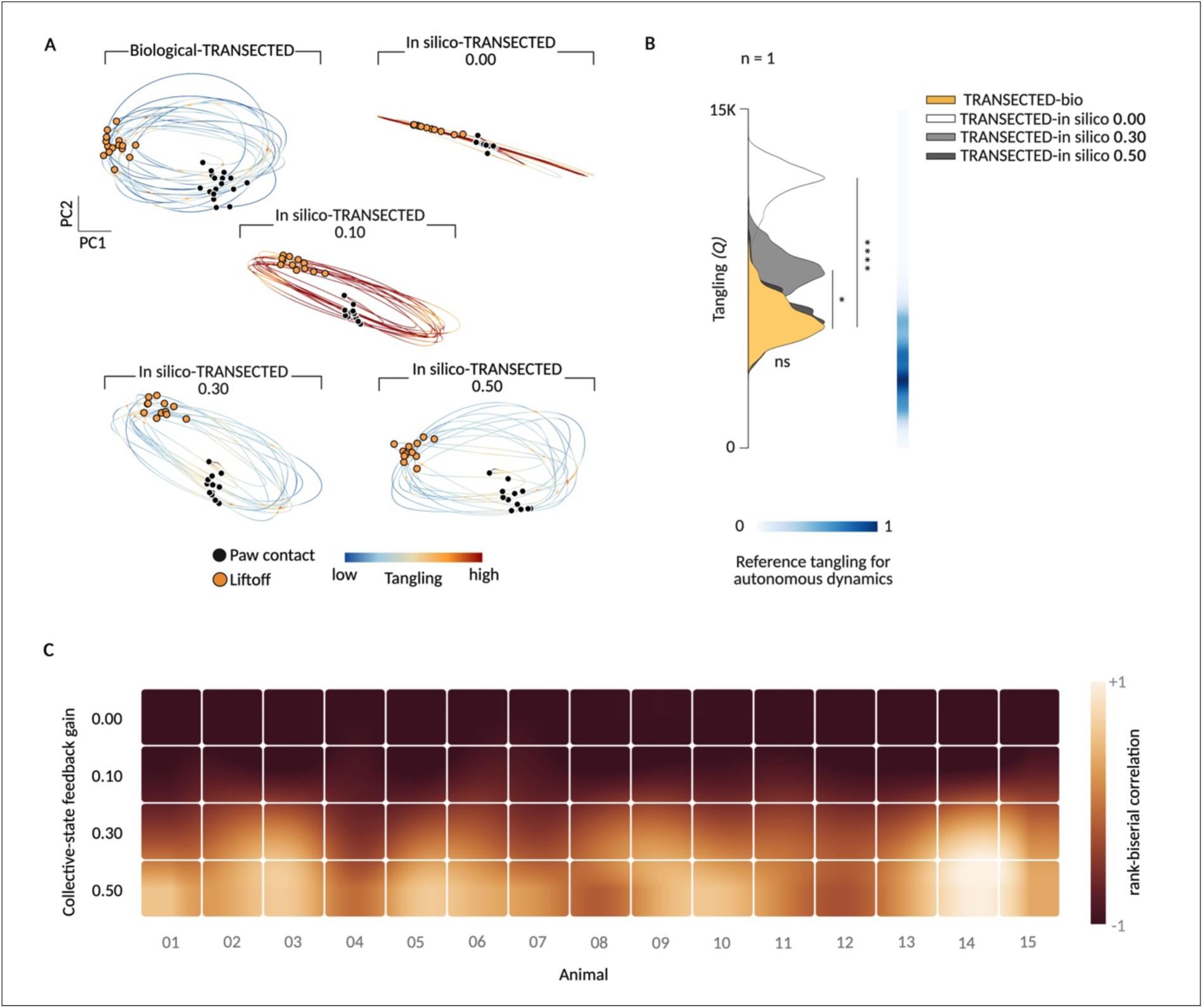
Intrinsic low-dimensional collective state dynamics organize low-tangled trajectories. **A)** Motoneuronal trajectories projected onto the PC1–PC2 plane for biological data (Biological-TRANSECTED) and in silico models with different collective-state feedback gains (0.00, 0.10, 0.30, and 0.50). Trajectories are colour-coded according to tangling values (low, blue; high, red). Black and orange dots indicate paw contact and liftoff, respectively. **B)** Quantification of tangling across biological and computational conditions in a representative animal. Violin plots show the distribution of cycle-wise root-mean-square tangling values. ****, P < 0.0001; *, P < 0.05; ns, not significant. **C)** Per-animal comparison of trajectory tangling between the biological transected condition and each collective-state feedback gain. Colour encodes the rank-biserial correlation of a two-sided Wilcoxon rank-sum test computed separately for each cat (n = 15). Negative values indicate lower tangling in the biological condition and the scale is centred on zero. Statistics for all 60 comparisons are in **Supplementary Table 11**.

## DISCUSSION

Our findings challenge the canonical framework of spinal locomotor control. Rather than arising from alternating flexor and extensor modules driven by somatosensory feedback^2,3,12,15^, locomotion reflects a low-dimensional dynamical process unfolding within spinal networks, taking the form of a rotation in neural space, as previously demonstrated in turtle rhythmic movements ^5^ and in intact rat locomotion^16^. During locomotion following spinal transection, the activity of the spinal motoneuron population continuously cycles through all phases, consistent with a limit-cycle attractor^17,18^, whereas the resulting global EMG activity remains alternating **(Fig. 1)**.

Building on dynamical systems approaches developed in the motor cortex^19,20^, we find that the rotational structure and its topology remain qualitatively preserved, although quantitatively degraded, despite the absence of supraspinal input, indicating that locomotor dynamics are intrinsically generated within the spinal circuitry **(Fig. 3)**. These low-dimensional, limit-cycle dynamics are expressed at the level of motoneuron populations, demonstrating that motoneurons are active readouts of premotor interneuronal circuits, dynamically co-varying with the underlying interneuronal network. This perspective is directly supported by recent recordings establishing a high-fidelity mapping (R^2^ > 0.83) and millisecond-scale temporal precision between spinal interneuronal state space and muscle activity^13^. While motoneurons function as active participants in the network’s collective state, the generation and organisation of these fundamental spinal dynamics are likely dominated by premotor interneuronal circuits.

Spinal transection does not abolish locomotion: spinal circuits remain capable of generating rhythmic activity in the absence of supraspinal input, but flexibility and adaptability are reduced^2,3,12^. Population activity becomes confined to a subspace with fewer modes, resulting in decreased effective dimensionality and trajectories that are less precisely organised. This shift likely reflects the loss of descending modulation of network gain acting on the spinal “projectome”, defined as the structured set of recurrent projections linking neuronal populations. The geometry of this projectome shapes the flow of activity in state space, consistent with theoretical work on recurrent attractor networks^21^ and recent studies of spinal population dynamics^22^. In intact conditions, supraspinal inputs modulate the gain of specific interneuron subpopulations^23^, thereby tuning the excitation/inhibition balance and shaping the geometry of the underlying projectome. This enables flexible control of the flow in state space and transitions between dynamical regimes. In their absence, the recurrent architecture continues to sustain rotational dynamics, but in a low-gain regime in which trajectories are less robust, less diverse, and less precisely aligned with the intrinsic flow field. This framework indicates that locomotion cannot be understood as a purely input-driven process. Feedforward extensions of classical modular schemes generate frequency-dependent phase shifts that are incompatible with stable coordination. Although somatosensory feedback can partially compensate for this, broad phase distributions persist even in somatosensory-deprived conditions^5,24,25^, indicating an additional intrinsic mechanism.

Using a recurrent neural network model constrained by biological data, we show that somatosensory feedback alone is insufficient to generate or sustain cyclic population dynamics. Instead, stable rhythmic activity emerges only when recurrent interactions are present, indicating that recurrent feedback of population activity is required **(Fig. 4 and 5)**. This interpretation is supported by the observation that slow rhythmic activity is consistently decoupled from fast synaptic potentials. Because input-driven control would predict strong temporal correlations across the population, this decoupling argues against feedforward-dominated models and instead supports a regime in which activity is shaped by internally generated dynamics^26^. A plausible explanation is that spinal circuits rely on recurrent, excitation/inhibition-balanced connectivity that stabilises rotational dynamics through internal feedback^27,28^.

In this context, somatosensory afferents act as modulators rather than primary drivers. They provide the input required to bring the system across a stability threshold and enable self-sustained activity, consistent with previous observations of intrinsic spinal dynamics^5^. This view complements models that emphasise a strong contribution of somatosensory feedback to locomotor control^9,29^.

This distinction is further supported by trajectory tangling analysis **(Fig. 6)**, which shows that, in the absence of intrinsic structure, population activity becomes highly sensitive to perturbations and weakly constrained by its own state. Restoring collective-state feedback recovers low-tangling values consistent with stable, locally predictable flow, thereby reinforcing the interpretation of locomotion as a self-sustained dynamical process.

Taken together, these findings support a fundamental revision of locomotor control. Our findings are consistent with a model in which locomotor rhythm is not imposed by descending commands or reconstructed from somatosensory feedback but instead reflects the intrinsic dynamics of spinal networks. Within this framework, supraspinal and sensory inputs do not generate locomotion. They shape, stabilize, and constrain the network states from which locomotor behaviour emerges.

This perspective provides a unifying account of both physiological function and dysfunction. Spinal cord injury may not eliminate the locomotor generator itself but instead compress the dynamical landscape available to spinal circuits, reducing their capacity to access and maintain functional network states. Recovery may therefore depend less on reconstructing neural pathways than on restoring the ability of spinal networks to explore and stabilize appropriate dynamical regimes.

The implications extend to current neuromodulation strategies. Although dorsal spinal cord epidural stimulation can restore stepping through the recruitment of sensory pathways, the resulting locomotion often remains constrained, less fluid, and only partially functional. Our findings suggest that future interventions should seek not to impose movement onto the spinal cord, but to re-enable the network mechanisms from which locomotion naturally emerges. Rather than driving the system from the outside, the goal may be to restore the conditions that allow spinal circuits to generate, sustain, and adapt their own locomotor dynamics.

Several limitations should be noted. First, motor units were sampled from eight muscle pools with fixed burst phases, so the present data cannot exclude that part of the observed phase structure reflects muscle identity rather than distributed population coding. Second, intact animals walked quadrupedally whereas transected animals stepped with the hindlimbs only, so differences in dimensionality and trajectory stability may partly reflect the change in task rather than the loss of supraspinal drive alone. Third, the model was trained exclusively on intact locomotion and transection was simulated by input ablation, which does not capture the circuit reorganisation occurring over the weeks following injury. Finally, because the model was trained with the collective-state feedback active, ablating it places the network outside its training distribution. The resulting loss of cyclic structure should therefore be interpreted as consistent with, rather than proof of, a structural requirement for output feedback.

## MATERIAL AND METHOD

### Animals and ethical information

All procedures were approved by the Animal Care Committee of the Université de Sherbrooke and were used in accordance with policies and directives of the Canadian Council on Animal Care (Protocol 2022–3349). Data from fifteen adult domestic short-haired cats (9 females and 6 males, >1 year of age at the time of experimentation) were used in the present study. We followed ARRIVE guidelines for animal studies^30^. Before and after experiments, cats were housed and fed in a dedicated room within the animal care facility of the Faculty of Medicine and Health Sciences at the Université de Sherbrooke. To reduce the number of animals used in research, cats participated in other studies to answer different scientific questions, many of which have been published^29,31–40^.

### Surgical procedures for electrode implantation

We performed surgeries under aseptic conditions with sterilised instruments in an operating room. Prior to surgery, the cat was sedated with an intramuscular injection of a cocktail containing butorphanol (0.4 mg/kg), acepromazine (0.1 mg/kg) and glycopyrrolate (0.01 mg/kg) and inducted with another intramuscular injection (0.05 ml/kg) of ketamine (2.0 mg/kg) and diazepam (0.25 mg/kg) in a 1:1 ratio. Cats were then anaesthetised with isoflurane (1.5–3%) and O_2_ using a mask for a minimum of 5 min and then intubated with a flexible endotracheal tube. Anaesthesia was maintained by adjusting isoflurane concentration as needed and by monitoring cardiac and respiratory rates. Body temperature was monitored with a rectal thermometer and maintained within physiological range (37 ± 0.5°C) using a water-filled heating pad placed under the animal and an infrared lamp ∼50 cm over it. We confirmed the depth of anaesthesia by applying pressure to a paw (to detect limb withdrawal) and by assessing the size and reactivity of pupils. The animal’s skin was carefully shaved using electric clippers and cleaned with chlorhexidine soap. We inserted a 24-gauge catheter in the left or right cephalic vein to give cats a continuous infusion of lactated Ringer’s solution (3 ml/kg/h). At the end of surgery, we injected an antibiotic (Convenia, 0.1 ml/kg) subcutaneously and taped a transdermal fentanyl patch (25 μg/h) to the back of the animal 2–3 cm rostral to the base of the tail for prolonged analgesia. We also injected buprenorphine (0.01 mg/kg), a fast-acting analgesic, subcutaneously at the end of the surgery and ∼7 h later. After surgery, we placed the cat in an incubator until it regained consciousness. We removed the fentanyl patch 4–5 days after surgery. At the conclusion of the experiments, cats received a lethal dose of pentobarbital through the cephalic vein. To confirm complete spinal transection in all cats, we performed histological analyses, a subset of which was previously reported in^33^.

### Electrode implantation

Cats were implanted with electrodes to chronically record the activity (EMG, electromyography) of several muscles. We directed pairs of Teflon-insulated multistranded fine wires (AS633; Cooner Wire, Chatsworth, CA, USA) subcutaneously from two head-mounted 34-pin connectors (Omnetics, Minneapolis, MN, USA). The two wires of each pair were twisted together along their length, crimped into a 21-gauge (0.8 mm) intramuscular needle used for insertion, and knotted together on the side opposite to the needle. Recording contacts consisted of 1–2 mm of insulation stripped from each wire between the knot and the needle, closer to the knot. Electrodes were inserted into the belly of selected hindlimb muscles for bipolar recordings. After insertion, the needle and excess distal wire were cut, and a distal knot was tied to anchor the electrode within the muscle. The head connector was secured to the skull using dental acrylic and six metallic screws.

### Spinal cord transection

The skin was incised over the 12th and 13th thoracic vertebrae, and after setting aside muscle and connective tissue, a small laminectomy of the dorsal bone was made. After exposing the spinal cord, we applied xylocaine (lidocaine hydrochloride, 2%) topically and made two to three injections within the spinal cord. We then completely transected the spinal cord with surgical scissors. We then cleaned the ∼0.5 cm gap between the two cut ends of the spinal cord and stopped any residual bleeding. We verified that no spinal cord tissue remained connecting rostral and caudal ends, which we later confirmed histologically. A haemostatic material (Spongostan, CDMV, Saint-Hyacinthe, QC, CAN) was inserted within the gap, and muscles and skin were sewn back to close the opening in anatomical layers. After spinal transection, we manually expressed the cat’s bladder and large intestine 2–3 times daily. Cats were then monitored daily by experienced personnel. The hindlimbs were cleaned as needed to prevent infection.

### Experimental protocol and data collection

We collected data (EMG and kinematics) before (intact state) and after spinal transection (transected state) during forward locomotion on a split-belt treadmill, with the left and right sides on separate belts. In the intact state, cats performed quadrupedal locomotion, while in the transected state, they performed hindlimb-only locomotion with the forelimbs on a stationary platform. During tied-belt locomotion, both sides stepped at 0.4 m/s. We collected data from 25-30 consecutive step cycles. In the transected state, we collected data between the 6th and 8th week after spinal transection when cats had a robust walking pattern.

We collected kinematic data by capturing videos of the left and right sides using two cameras (Basler AcA640-100 g, Basler AG, Ahrensburg, Germany) at 60 frames/s with a spatial resolution of 640 × 480 pixels. A custom-made program (Labview, National Instruments, Austin, TX, USA) acquired the images and synchronised acquisition with EMG data. By visual detection, we determined limb contact as the first frame where the paw made visible contact with the treadmill surface, and limb liftoff as the most caudal displacement of the toe, for both hindlimbs.

EMG signals were pre-amplified (10×, custom-made system), bandpass filtered (30– 1000 Hz) and amplified (100–5000×) using a 16-channel amplifier (model 3500; A-M Systems, Sequim, WA, USA). EMG data were digitized (2000 Hz) with a National Instruments card (NI 6032E), acquired with custom-made acquisition software and stored on computer. We analysed the following muscles: medial and lateral gastrocnemius (MG, LG, ankle extensors/knee flexors), soleus (SOL, ankle extensor), tibialis anterior (TA, ankle flexor), biceps femoris posterior (BFP, knee flexor/hip extensor), vastus lateralis (VL, knee extensor), anterior sartorius (SRT, hip flexor/knee extensor) and semitendinosus (ST, hip extensor/knee flexor). Although some muscles have both extensor and flexor actions, we functionally define extensors as those mostly active during stance (MG, LG, SOL and VL) and flexors as those mostly active during swing (TA, BFP, SRT and ST).

### Decomposition of EMG signal

Motor unit analysis was performed offline in Spike2 (version 10; Cambridge Electronic Design, Cambridge, UK), and spike rasters were aligned to the gait cycle using a custom Python-based code. The full decomposition pipeline is illustrated in **Supplementary Figure 1**. EMG signals were high-pass filtered on each channel using a second-order Butterworth filter (200 Hz cutoff), and motor unit spike times were identified using voltage threshold detection followed by whole-waveform template matching, as previously described for fine-wire recordings during locomotion^14^. Templates were allowed to track slow changes in waveform shape over a recording, and highly similar templates were merged. Sorting was verified in the cluster window, where spikes were projected onto the first three principal components and separated using the normal-mixtures algorithm. Units were then manually curated. Units were accepted based on cluster separability, waveform consistency, and a low incidence of refractory-period violations in the inter-spike interval histogram^41,42^. Motor unit spike trains were finally segmented according to locomotor phase using externally recorded gait event markers, enabling phase-specific analyses of motor unit activity.

### Euthanasia and histology

At the end of the experiments, cats were anaesthetised with isoflurane (1.5–3.0%) and O2 before receiving a lethal dose (100 mg/kg) of pentobarbital through the left or right cephalic vein. Using a stethoscope, we confirmed cardiac arrest. To confirm that spinal transection was complete in all cats, we performed a visual inspection during the surgery and a post-mortem histological analysis^33^. Briefly, we removed a 2 cm-long segment of the spinal cord around the spinal lesion immediately after euthanasia. We then placed the spinal segment in 25 ml of a 4% paraformaldehyde solution in 0.1 M phosphate-buffered saline (PBS) solution at 4°C. After 5 days, the spinal segment was cryoprotected in PBS with 30% sucrose for 72 h at 4°C. Coronal sections of 50 μm of the spinal cord were mounted on slides and stained with 1% cresyl violet. We then performed qualitative and quantitative evaluations of the injury site.

## ANALYSES

### Motor-unit analysis

All data analyses were performed using custom-designed procedures in Python (version 3.13). Motor-unit spike trains were synchronised to the locomotor cycle using kinematic events, including paw contact and lift-off. For visualisation and all subsequent population analyses, spike trains were binned at 2 ms resolution (500 Hz), square-root transformed to stabilise the variance of the spike-count process, and convolved with a Gaussian kernel^43^:

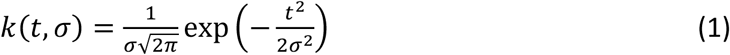

Where σ = 50 ms, yielding continuous estimates of motor-unit discharge rates throughout the locomotor cycle.

Two complementary representations of motor-unit activity were generated, both aligned to the locomotor cycle. First, spike rasters were displayed with motor units ordered according to their extraction sequence (chronological sorting). Second, both spike rasters and continuous discharge-rate estimates were reordered according to each unit’s preferred firing phase, defined as the circular mean of its discharge times across the locomotor cycle (phase sorting, or resorting).

Motor-unit discharge behaviour was quantified separately during stance and swing and compared between intact and transected conditions. For each identified motor unit, the mean discharge rate (pps) and discharge duration were calculated within each locomotor phase and averaged across valid locomotor cycles. Discharge duration was expressed as a percentage of phase duration to account for differences in cycle timing.

### Manifold-based analysis of population dynamics

For each experimental condition, the continuous discharge-rate estimates described above were concatenated across locomotor cycles to construct a population activity matrix X ∈ R^n×T^, where *n* represents the number of motor units and T is the total cumulative duration.

The manifold structure was identified using Principal Component Analysis (PCA). The principal components and their associated eigenvalues were determined as eigenvectors of the empirical covariance matrix. The eigenvectors and eigenvalues were found through the eigen-decomposition:

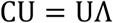

where U contains the principal components (eigenvectors), and **Λ** is a diagonal matrix containing the eigenvalues, representing the variance explained by each component:

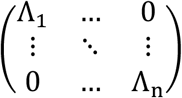

### Motoneuronal variance accounted for (VAF)

We used cumulative variance accounted for (VAF) analysis to determine the dimensionality of the manifold, defined as the minimum number of motoneuronal modes required to explain motor-unit population activity^44^. The representativeness of the manifold was quantified by the cumulative VAF (VAFh) for a subset of h modes:

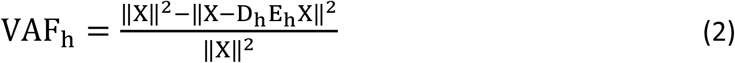

where X is the centered data matrix, E_h_ is the encoding matrix, and D_h_ is the decoding matrix.

### Canonical correlation analysis (CC)

CC was used to quantify the preservation of motor trajectories and dimensional spaces between different conditions (e.g., Intact vs. Transected). This method identifies directions within the respective manifolds where the latent activities are maximally correlated^45^. CC coefficients were obtained via singular value decomposition of the product of the orthonormal bases Q_A_and Q_B_ of the compared manifolds:

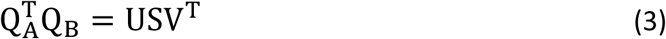

The diagonal elements of matrix S correspond to the CC coefficients. A global stability score was calculated by averaging the CCs of the first five motoneuronal modes. This extends the top four canonical correlation metric introduced by Gallego et al.^46^ by including a fifth mode, corresponding to the dimensionality required to explain 80% of the population variance in intact animals **(Fig. 3B)**, thereby ensuring that all modes contributing substantially to the manifold were retained. The significance of CC was assessed using a time-shuffling control procedure. Artificial surrogate datasets were generated by randomly permuting the time indices of the latent activity over the entire concatenated duration T. To preserve the spectral structure induced by the initial smoothing, these shuffled data were re-smoothed with the same 50 ms Gaussian kernel before being subjected to CC. This procedure was repeated 5000 times to generate a null distribution of canonical correlations. The significance threshold was set at the 99.9th percentile of the distribution of these random correlations (P < 0.001).

### Manifold Topology

To characterise the geometry of the trajectory manifold within each condition, we applied persistent homology to the low-dimensional neural trajectories. Persistent homology was computed on the first five principal components of the manifold identified above. The trajectories were subsequently subsampled to reduce computational complexity while preserving their geometric structure. Persistent homology was then computed using a Vietoris–Rips filtration implemented in *Ripser*. Formally, for a set of points *X* in the latent space and a radius ɛ, the Vietoris-Rips complex VR(X, ɛ) is defined as the set of all simplices σ ⊆ X such that all pairs of vertices in σ are within a distance ɛ.

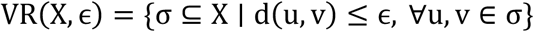

In this framework, H_0_ features correspond to connected components, H_1_ features capture loop-like structures reflecting cyclic organisation of trajectories, and H_2_ features indicate higher-dimensional cavities. The presence of H_1_ without H_2_ therefore indicates a solid ring– like topology rather than a toroidal surface, consistent with cyclic dynamics on a low-dimensional manifold. Persistence diagrams were visualised as barcode representations to highlight the most stable topological features.

### Distance to the mean cycle

To quantify the stability of the trajectory manifold across locomotor cycles, we measured the deviation of individual cycles from the mean trajectory in the latent state space. Locomotor cycles were then segmented based on detected gait events and resampled to a fixed number of phase points (N = 100) to normalise cycle duration. For each condition, an average cycle trajectory xT was computed across all normalised cycles. The stability of each individual cycle k was quantified as the mean Euclidean distance between the cycle trajectory x_k_ and the mean cycle across corresponding phase points.

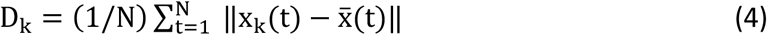

This metric, denoted D_k_, captures the dispersion of individual trajectories around the limit-cycle manifold, with lower values indicating more stable dynamics.

### Trajectory alignment and flow

To quantify the dynamical organisation of neural activity in state space, we estimated the trajectory alignment and the flow field within the low-dimensional manifold. Conceptually related to previous alignment analyses^16^, trajectory alignment (A) was defined as the normalised projection (cosine similarity) between the instantaneous velocity ẋ(t) and the local flow vector F(x(t)) at the corresponding location in state space.

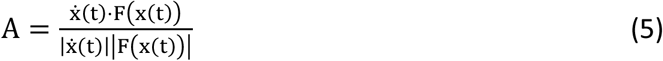

Alignment values were averaged across time and normalised relative to a shuffled baseline obtained by temporally permuting the trajectories. To determine the local flow vectors, a continuous flow field F(v) was estimated for each voxel v on a spatial grid by locally averaging instantaneous velocity vectors ẋ(t) using a Gaussian kernel

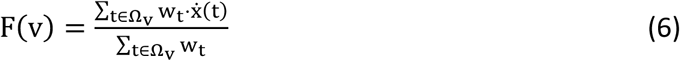

where Ω_v_ represents the set of points in the neighborhood of voxel v, and w_t_ is a Gaussian weight based on the distance to the voxel center. The state space was discretised into voxels, and trajectory occupancy was used to identify regions belonging to the manifold. Voxels repeatedly visited by the trajectories were defined as inside the manifold, whereas sparsely visited regions were considered outside the manifold. Attractor strength was then quantified by computing the mean flow magnitude separately for these two regions. High alignment and stronger flow within the manifold indicate coherent motion consistent with limit-cycle dynamics.

### Local Lyapunov exponents

To quantify the local stability of motoneuronal trajectories, we computed the Lyapunov exponent (λ). Following the framework of Komi et al.^16^, for each point in the latent state space, pairs of trajectories were identified via neighbor selection, and their temporal evolution was tracked over a short time horizon. The Lyapunov exponent was defined as the average logarithmic rate of divergence (or convergence) between these trajectory pairs:

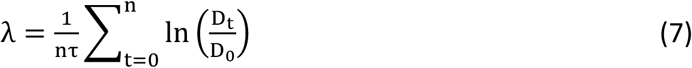

where n is the number of samples, D_0_ is the initial Euclidean distance between trajectories and D_t_ is the distance after the time interval τ.

In this framework, a near-zero exponent (λ ≈ 0) indicates that trajectories co-evolve and remain at a near-constant distance, characteristic of the stable limit cycle dynamics, λ < 0 indicates attractor dynamics where trajectories converge over time towards a stable state, such as the fixed points and λ > 0 indicates unstable dynamics where nearby trajectories rapidly separate.

### Tangling analysis of population dynamics

To determine whether model-generated population activity was governed primarily by intrinsic collective dynamics or by externally driven inputs, we quantified the local structure of neural trajectories using the trajectory tangling metric. Tangling assesses the extent to which the short-term evolution of a trajectory can be predicted from its instantaneous position in neural state space. Specifically, it measures whether nearby population states evolve with similar velocities, indicating locally consistent and predictable flow or with divergent velocities, indicating sensitivity to external inputs or perturbations.

This framework has been widely used to distinguish population dynamics that are predominantly governed by collective state dynamics from those strongly influenced by external drive^20,47^. Low tangling indicates that the future evolution of population activity is largely determined by its current state and is therefore locally predictable. In contrast, high tangling reflects divergence of future trajectories from nearby states, consistent with externally driven or perturbed dynamics. Importantly, tangling should be interpreted as a relative measure of local dynamical predictability rather than as an absolute classifier separating autonomous from input-driven systems.

Analyses were performed on continuous segments of total population activity (sampled at 500 Hz). Trajectory tangling was computed from the continuous discharge-rate estimates described in the Motor-unit analysis section. To ensure that comparisons between simulated and biological activity were not driven by trivial differences in global amplitude, simulated activity was additionally rescaled to match the 99th percentile of the absolute biological firing rates prior to dimensionality reduction.

Trajectory tangling was computed using the first five principal components of the manifold identified above, corresponding to the dimensionality required to explain approximately 80% of the population variance in intact animals. This yielded a five-dimensional neural state representation x(t) ∈ ℝ^5^ capturing the dominant population dynamics. Using a fixed latent dimensionality ensured consistent tangling estimation across datasets.

### Tangling metric

Tangling was computed from the instantaneous population state x(t) and its temporal derivative ẋ(t) which was estimated by numerical differentiation. For each time point t, tangling was defined as:

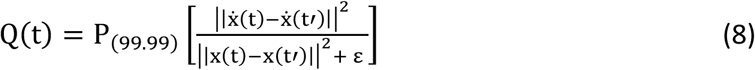

where *t*′ indexes all other time points, ε = 10⁻⁶ prevents division by zero and the 99.99^th^ percentile (P_99.99_) was used to obtain a robust estimate of maximal local divergence while minimising sensitivity to rare numerical outliers^48^. To evaluate the stability of functional locomotor units, data were segmented into individual gait cycles, and the root-mean-square of the instantaneous values was calculated for each cycle. This formulation follows that introduced by Russo et al.^47^ and subsequently applied to naturalistic behaviour by Perich et al.^48^, whereby tangling increases when nearby states in neural state space evolve with dissimilar velocities, consistent with externally driven or perturbed dynamics.

Tangling distributions were computed for the biological spinal-transected reference population, and in silico-transected model simulations generated under different collective-state feedback gains. Each data point in the resulting distributions represents the root-mean-square tangling of an individual gait cycle. These analyses were performed independently for each animal, and differences between conditions were statistically tested for each subject using the Wilcoxon rank-sum test. To provide a visual reference for dynamics primarily driven by collective-state feedback rather than by structured external inputs, we computed a reference tangling distribution using the RNN model operating in an autonomous regime. In this configuration, supraspinal drive and phase-dependent somatosensory inputs were suppressed, so that network activity evolved mainly from its own collective-state feedback. Under these conditions, population trajectories reflect self-generated collective state dynamics with minimal direct external drive. This autonomous RNN regime was used as a reference baseline for interpreting tangling levels across conditions.

### Latent recurrent neural network model

We developed a latent recurrent dynamical model fit to the combined population of motor units recorded across multiple muscles. Although motor unit decomposition was performed independently for each muscle, firing-rate vectors were constructed by concatenating motor units identified across muscles and temporally aligned to a common locomotor phase. As a result, the model operated on a single multi-muscle population vector at each time point, allowing the latent dynamics to capture distributed, population-level coordination spanning multiple muscles rather than modeling each muscle independently. The latent recurrent state was trained to reproduce the observed motor unit population activity, thereby providing a compact representation of the collective dynamics underlying locomotor output. Through training, the recurrent network learned the recurrent transformations linking structured inputs to motor unit population activity, enabling in silico perturbations to causally determine how specific inputs contribute to the emergence of locomotor dynamics. To investigate how rhythmic motor unit population activity emerges from the interaction between structured inputs and intrinsic spinal dynamics, we modeled motor unit discharge rates using a recurrent latent dynamical system. The observed activity is represented by:

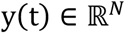

the vector of firing rates of N motor units at time t, obtained from intramuscular EMG decomposition. Firing rates were computed as described above (see Motor-unit characterization). To reduce amplitude differences across motor units and stabilise model training, each unit’s firing-rate trace was normalised by its maximum value over the recording. The model assumes that observed population activity arises from a low-dimensional latent state

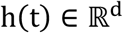

The latent dimensionality d corresponds to the number of recurrent units per layer in a two-layer Gated Recurrent Unit (GRU) architecture (128 units per layer). This dimensionality provided sufficient capacity to capture collective recurrent population dynamics while remaining low-dimensional relative to the number of observed motor units (N). The temporal evolution of the latent state was governed by a causal recurrent update rule implemented as a GRU:

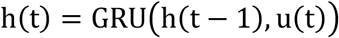

Formally, this update comprises a reset gate and an update gate that jointly regulate the integration of new input relative to the previous hidden state:

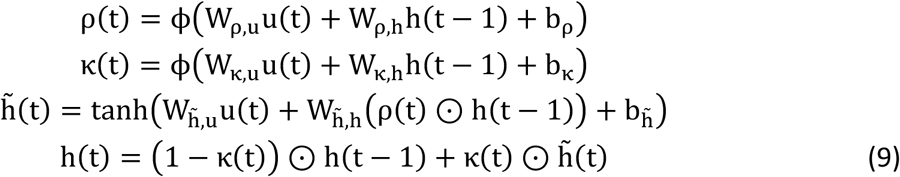

where **φ**(⋅) denotes the logistic sigmoid function, ⊙ the element-wise product, and W and b the learned parameters of the recurrent layer. The network was implemented in PyTorch, whose GRU is equivalent to this formulation up to a relabelling of the update gate.

Where h(t) ∈ ℝ^H^denotes the latent population state at time t, u(t) is a low-dimensional input vector that integrates structured locomotor signals together with a compressed representation of recent population activity. This recurrent formulation enforces temporal causality and incorporates multiplicative gating mechanisms that regulate information flow across time, allowing the latent state to integrate inputs over multiple timescales while maintaining stable recurrent dynamics^49,50^.

The input vector u(t) comprised a small set of low-dimensional signals capturing the temporal structure of locomotion rather than unit-specific tuning. These included: **(i)** a constant bias term, interpreted as a descending supraspinal drive and **(ii)** a phase-like signal that increased monotonically within each gait cycle, resetting at cycle boundaries to serve as a proxy for phase-dependent somatosensory feedback.

### Collective-state feedback input

To provide the recurrent network with access to recurrent collective population activity while avoiding direct high-dimensional feedback, we introduced a low-dimensional collective state dynamics signal derived from the previous firing-rate vector y(t − 1). At each time point, firing rates were first centered by subtracting the instantaneous population mean, thereby removing global fluctuations shared across motor units. The centered population vector was then projected into a 12-dimensional collective state representation using a compact learned encoder (two fully connected layers with 64 hidden units and a hyperbolic tangent nonlinearity, followed by layer normalisation), trained jointly with the recurrent network via backpropagation rather than fitted as a separate unsupervised step:

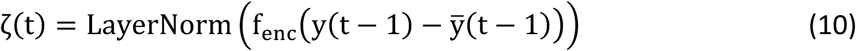

Because this encoder is fully differentiable, gradients propagate through the reinjected collective state signal itself, not only through the recurrent hidden state, allowing the network to learn a representation directly optimised for stable, self-consistent generation.

The resulting low-dimensional state representation vector ζ(t) was concatenated with the structured locomotor inputs to form the complete GRU input u(t). This procedure yields a low-dimensional representation of recent population dynamics that stabilises training and encourages the latent state to capture structured collective dynamics rather than unit-specific noise. Importantly, this low-dimensional state signal is not an independent external input but rather an explicit representation of the system’s collective dynamics. This formulation enables controlled manipulation of intrinsic population activity, allowing us to directly probe the contribution of intrinsic network dynamics to the emergence of rhythmic locomotor patterns. The recurrent latent state was layer-normalised and mapped to motor unit activity by a linear readout with a separate bias, followed by a softplus nonlinearity to ensure non-negative firing rates.

### Training procedure and held-out evaluation

Model parameters were optimised using temporally segmented data to prevent information leakage between training and evaluation sets. Continuous recordings were split along the time axis into non-overlapping segments, with 80% of the data used for training and the remaining 20% held out for evaluation. All reported performance metrics were computed exclusively on the held-out data.

The model was optimised using truncated backpropagation through time over non-overlapping segments of 250-time bins, combined with scheduled sampling. At each time step within a segment, the low-dimensional collective state signal was derived either from ground-truth motor-unit activity or from the network’s own previously generated output, with the probability of using ground-truth activity decreasing linearly over training (from 1.0 to 0.2)^51^.

For held-out evaluation, this signal was instead constructed exclusively from ground-truth activity. Model performance was quantified using the coefficient of determination (R^2^), computed separately for each motor unit by comparing predicted and observed firing rates on held-out data. Population-level performance was summarised by averaging R^2^ values across units, excluding undefined values when variance was insufficient.

### Reliability analysis and noise ceiling estimation

To estimate the upper bound on explainable variance for each motor unit, we computed a reliability-based noise ceiling using a split-half procedure across gait cycles. For each unit, firing-rate profiles were extracted on a cycle-by-cycle basis, resampled to 100 phase points, and averaged separately across odd and even cycles. The Pearson correlation between these two averaged profiles was corrected using the Spearman–Brown formula and squared to express the ceiling as a fraction of explainable variance comparable to R².

This reliability estimate provides an upper bound on the fraction of variance that can be explained by any model, given the intrinsic trial-to-trial variability of the motor-unit activity. Motor units were retained for subsequent analyses if held-out R² exceeded 0.4 or reached at least 60% of the estimated noise ceiling for units whose ceiling exceeded 0.1, ensuring that units with low apparent performance due to limited reliability were not spuriously excluded. This approach distinguishes model limitations from intrinsic variability in motor-unit firing, providing a principled criterion for unit inclusion.

### Input ablation and sufficiency analyses

In all configurations, the recurrent update of the GRU remained unchanged. The manipulations described below act exclusively on the input vector u(t), including its collective-state feedback component ζ(t). To assess the relative contribution of structured external inputs and collective-state feedback to the learned population activity, we performed a series of controlled input-ablation and sufficiency analyses. Model predictions were generated by selectively suppressing specific components of the input vector u(t) at inference time, while keeping all network parameters fixed. Critically, these predictions were generated in a purely autoregressive, closed-loop manner: following a brief initialisation period (250-time bins) during which the collective state signal was derived from ground-truth activity to establish a physiologically plausible hidden state, all subsequent time steps used the collective state signal derived exclusively from the network’s own previously generated output, with no further access to ground-truth activity. This ensured that the resulting trajectories reflected the network’s intrinsic capacity to sustain structured activity under each input configuration, rather than activity partially reconstructed from held-out biological recordings.

We defined the following experimental conditions. **In silico-intact** used the full input vector, including supraspinal drive and phase-dependent somatosensory input, and low-dimensional collective state components. **In silico-autonomous** suppressed all structured external inputs (supraspinal and somatosensory), while maintaining low-dimensional collective state dynamics at full strength. **In silico-transected** suppressed supraspinal drive while preserving somatosensory input and low-dimensional collective state dynamics. In silico-somatosensory-only retained only the phase-related somatosensory input while suppressing the supraspinal bias and the collective-state feedback, testing the sufficiency of somatosensory phase information alone.

### External validation against an independent population reference

To evaluate whether spinal-only model predictions captured biologically meaningful population dynamics, we performed an external validation against an independent biological reference dataset recorded in the same cat before and after spinal transection. In the model, supraspinal inputs were explicitly removed, such that simulated motor-unit activity reflected population dynamics generated solely by intrinsic spinal circuitry.

Rather than treating internal recurrence as an all-or-none mechanism, we systematically modulated the contribution of the collective-state feedback. Specifically, the gain applied to the low-dimensional collective state signal (i.e., itself derived from the network’s own previously generated activity under the closed-loop generation procedure) was scaled, allowing its contribution to be progressively increased from zero to full strength. This manipulation enabled a direct test of the necessity of collective-state feedback, as opposed to residual ground-truth information, to sustain structured population activity in the absence of supraspinal drive. Model simulations generated under these spinal-only conditions were compared to biological motor-unit recordings obtained after spinal transection in the same animal, enabling a within-subject comparison under matched functional conditions. The biological reference dataset was processed using the same preprocessing pipeline as the intact recordings used to train the model, including temporal binning, smoothing and normalisation, ensuring methodological consistency across intact, simulated, and post-spinal datasets.

Because motor units were not in one-to-one correspondence between model outputs and biological recordings, a data-driven matching and evaluation procedure was employed. To avoid circularity, the post-spinal transection biological dataset was split into two non-overlapping temporal segments of equal duration. The first segment (50% of the data) was used exclusively to estimate unit correspondence based on covariance structure and to define a low-dimensional reference population manifold using PCA. No information from the remaining segment was used during this stage.

Model predictions and biological activity from the remaining held-out segment (the other 50% of the data) were then projected into this fixed reference manifold and used solely for evaluation. To account for small temporal misalignments, an optimal lag was estimated using cross-correlation prior to computing similarity. Additionally, component polarity was aligned to account for the arbitrary sign of PCA axes. By evaluating model–biological similarity exclusively on held-out data as a function of recurrent latent-state strength, this analysis provided a stringent external test of whether intrinsic spinal recurrent dynamics were necessary to reproduce the population-level rhythmic motor-unit activity observed following spinal transection.

### Statistical analysis

Statistical analyses were performed in R (version 4.3.2). Computational simulations and post-processing were conducted in Python (version 3.13) within Jupyter notebooks. Graphs were generated using Python and GraphPad Prism (version 10.0; GraphPad Software, USA), and schematic illustrations were created with Python and BioRender. Statistical significance was set at P < 0.05 for all analyses.

### Experimental analysis

Experimental data were independently collected from 15 adult cats across multiple sessions. For each animal and each experimental condition (Intact and Transected), a sequence of 25 to 30 consecutive gait cycles was analysed. Circular uniformity of motor unit discharge phases across the locomotor cycle was assessed independently for each animal using a Kuiper test. To establish a statistical standard, the same Kuiper test was applied to uniformly distributed surrogate datasets. The resulting experimental p-values (n = 15) were then compared to the null distribution of surrogate p-values using a non-parametric Wilcoxon rank-sum test. This approach accounted for the non-normal and bounded nature of p-values, providing a robust group-level assessment of circular uniformity by benchmarking experimental results against a theoretical null. All subsequent comparisons concerning motor unit characteristics, manifold geometry, and dynamical parameters were performed using Linear Mixed-Effects Models (LMM) with animal ID included as a random effect (random intercept) to control for inter-animal variability and the non-independence data from the same animal. For the analysis of motor unit discharge patterns, Condition (Intact vs. Transected), Phase (Stance vs. Swing), and Functional Group (Extensors vs. Flexors) were treated as fixed effects, including their interactions, with mean discharge frequency and normalised discharge duration as dependent variables. This LMM framework was similarly employed to evaluate global manifold and dynamical features, including number of dimensions to explain 80% of variance, distance to the mean cycle, and the Lyapunov exponent, between experimental conditions. Finally, trajectory alignment (comparing Intact vs. Transected and Original vs. Shuffled) and mean flow (comparing Intact vs. Transected and In-manifold vs. Out-manifold) were assessed using the same linear mixed-effects modeling approach.

### Simulated data analysis

Statistical validation of the computational models and specific dynamical features was performed using a series of LMM and non-parametric tests. To compare biological and simulated states, LMMs were constructed with Condition (biological Intact and models parameters) as a fixed effect and Cat ID as a random effect to account for inter-animal variability. This mixed-modeling approach was similarly applied to assess trajectory alignment and vector field flow, comparing conditions (Intact vs. Transected), data structures (Original vs. Shuffled) for alignment, and (In-manifold vs. Out-manifold) for mean flow. The correspondence between biological-intact population dynamics and the spinal in silico model with low-dimensional representation of collective state scaling was quantified by calculating the Pearson correlation coefficient between the respective PC1. Within each animal, trajectory tangling was compared between the Biological-TRANSECTED condition and each in silico condition (0.00, 0.10, 0.30, 0.50) using two-sided Wilcoxon rank-sum tests on per-gait-cycle values (*n* = 15 cats; 25–30 cycles per animal and condition). The four *P* values per animal were Holm–Bonferroni corrected and adjusted values are reported (descriptive statistics in **Supplementary Table 12)**. Effect sizes are given as the rank-biserial correlation, negative when tangling is lower in the biological condition.

## Data availability

All data are available from the corresponding author (A.F.) upon reasonable request and will be made publicly available in an open-access repository upon publication.

## Code availability

Code for the latent recurrent model and the in silico manipulations are available at https://github.com/JulianColard/Spinal-Locomotor-RNN.

## Acknowledgements

We thank students and staff in the Frigon lab that participated in data collection over the years. This work was supported by a Network Grant from the Collaboration on motor planning, execution and resilience (COMPERE).

## Authors contributions

J.C., J.H. and A.F. conceived the project. J.C. and J.H. performed the surgeries and experiments and collected the data. J.C., R.W.B. and A.F. conceived the original theory. J.C. and R.W.B. performed the model simulations and developed the population-dynamics analyses. J.C., J.H., R.W.B. and A.F. wrote the manuscript. A.F. supervised the project and acquired funding. All authors reviewed and approved the final manuscript.

## Competing interests

The authors declare that no competing interests exist.

## Notes

### Competing Interest Statement

The authors have declared no competing interest.

### Summary of Updates

Figure 6 has been revised, and an additional panel (C) has been added.

